# CNAttention: an attention-based deep multiple-instance method for uncovering CNA signatures across cancers

**DOI:** 10.1101/2025.05.26.656180

**Authors:** Ziying Yang, Michael Baudis

**Author notes:** Contributing authors.

## Abstract

Somatic copy number aberrations (CNAs) represent a distinct class of genomic mutations associated with oncogenetic effects. Over the past three decades, significant volumes of CNA data have been generated through molecular-cytogenetic and genome sequencing-based techniques. This data has been pivotal in identifying cancer-related genes and advancing research on the relationship between CNAs and histopathologically defined cancer types. However, comprehensive studies of CNA landscapes and disease parameters are challenging due to the vast diagnostic and genomic heterogeneity encountered in “pan-cancer” approaches. In this study, we introduce *CNAttention*, an attention-based deep multiple instance learning method designed to comprehensively analyze CNAs across different cancers and uncover specific CNA patterns within integrated gene-level CNA profiles of 30 cancer types. CNAttention effectively learns CNA features unique to each cancer type and generates CNA signatures for 30 cancer types using attention mechanisms, highlighting the distinctiveness of their CNA landscapes. CNAttention demonstrates high accuracy and exhibits stable performance even with the incorporation of external datasets or parameter adjustments, underscoring its effectiveness in tumor identification. Expanding these signatures to cancer classification trees reveals common patterns not only among physiologically related cancer types but also among clinico-pathologically distant types, such as different cancers originating from neural crest derived cells. Additionally, detected signatures also uncover genomic heterogeneity in individual cancer types, for instance in brain lower-grade glioma. Additional experiments with classification models underscore the efficacy of these signatures in representing various cancer types and their potential utility in clinical diagnosis.

## 1 Introduction

Copy number variations (CNVs) refer to changes in the number of copies of specific DNA segments in the genome, usually including deletions or duplications with sizes ranging from 1kp to multiple megabases [1]. Copy number aberrations (CNAs) refer to acquired CNVs in the disease genome in comparison to the healthy genome, potentially altering the diploid status of specific genomic loci. For instance, neurodevelopmental disorders like intellectual disability, autism, and schizophrenia have been found to be associated with CNAs, accounting for at least 15% of these conditions [2]. The relevance between CNAs and neurodevelopmental diseases may stem from the disruption of gene pathways involved in neuron development. Additionally, several genes implicated in neurodevelopment, such as A2BP1, IMMP2L, and AUTS2, have been reported to harbor mutational CNAs [3].

Cancer initiation and progression are closely linked to changes in copy number [4]. In breast cancer, for instance, Li et al. conducted an oncogenetic tree analysis of CNAs and found that the genetic alteration of ErbB2 occurs early, while CNAs of AKT2, PTEN, CCND1, RAS, and PIK3CA are late events [5]. This association can be partially attributed to cellular stress, as copy number changes often occur in response to stressors like hypoxia, which may switch DNA repair mechanisms from homologous recombination to non-homologous repair [6]. There is ample evidence suggesting that individuals with certain CNAs may be predisposed to cancer [7]. Despite this, the majority of studies have focused on identifying associations with cancer-driver genes or the impact of focal regions in specific tumor types. Consequently, CNA patterns have often been characterized by the coverage of driver genes, rather than through comparative analyses of the entire genome. However, this approach has two drawbacks. Firstly, the distribution of cancer driver genes is highly skewed, with only a few hallmark drivers accounting for a large percentage of tumorigenesis, leaving a long-tail of rare or putative drivers to account for the rest [8]. Secondly, research has revealed various facets of CNAs in relation to cellular regulations and genome dynamics [9– 12]. Therefore, CNA patterns that solely rely on driver genes often fail to capture the full spectrum of aberrations. To address this limitation, it would be more comprehensive to abstract CNA patterns based on their characteristic aberrations, rather than focusing solely on focal regions overlapping with driver genes.

While on a global scale, the associations between CNAs and different types of cancer still remain elusive. Some CNAs overlapped between tumor types, others were tumor type-specific; losses of CDH20 and PTEN were observed in both tumor types, whereas amplifications seemed more tumor type-specific, such as EGFR and MAP2K4 in colon cancer and ERBB2 in breast cancer [13]. Some previous studies have been able to delineate diverging patterns in clinically related entities. For example, it could be shown that the CNA patterns between lung adenocarcinoma and squamous cell carcinoma are very different [14] and that there are specific landscapes of mutations and copy number changes in various cancers [15]. Interestingly, cancer type-specific CNAs derived from cell-free DNA (cfDNA) have demonstrated their potential in identifying cancer types and tissues of origin [16],[17],[18]. As an emerging field, the accurate identification of genomic abnormalities and classifications of the cfDNA remains challenging, and therefore, the characterization of specific CNA patterns related to cancer types could provide valuable information for such applications.

In this study, we assembled a collection of 10,628 CNA profiles and introduced a novel attention-based method CNAttention, which by capturing specific weighted features of each cancer type in our analysis achieved a classification accuracy of 0.89. When testing our method on additional datasets the accuracy did not deteriorate, showcasing the effectiveness of CNAttention in extracting relevant CNA characteristics of different cancers and in identifying tumors types. Additionally, we used the weights assigned by the attention mechanism to generate specific CNA signatures for 30 cancer types and compared these signatures with the original data and indicators for their biological significance. The result illustrates the genetic uniqueness and relationships of extracted CNA patterns in different cancer types but also uncovers heterogeneity within cancer classifications, therefore demonstrating the potential of CNA signatures in improving diagnostic assessments in oncology.

## 2 Methodology

As shown in Fig 1, our method CNAttention is a three-stage method. Based on a training set it firstly applies RFECV to select important features, which remain the characteristics of CNA patterns of different cancer types. This step is for reducing the computational time and avoiding over-fitting caused by the much larger number of features compared to the number of samples. Next, we transfer the classification problem into a multiple-instance learning problem by importing the attention mechanism, to find out the key instances that better capture the CNA pattern of specific cancers. Lastly, by transferring the instance weights within bags to feature weights to cancer types, we generate the CNA signatures of each cancer type, which represents the CNA patterns across the cancer landscape. We randomly selected 80% of the gene-level CNA data from cBioPortal for training and reserved the remaining 20% for testing. To validate the CNA signatures on a larger CNV dataset, we mapped them to the Progenetix database (progenetix.org [19–21]), which curates a comprehensive reference collection of CNV-centered oncogenomic profiling data.

**Fig. 1.**
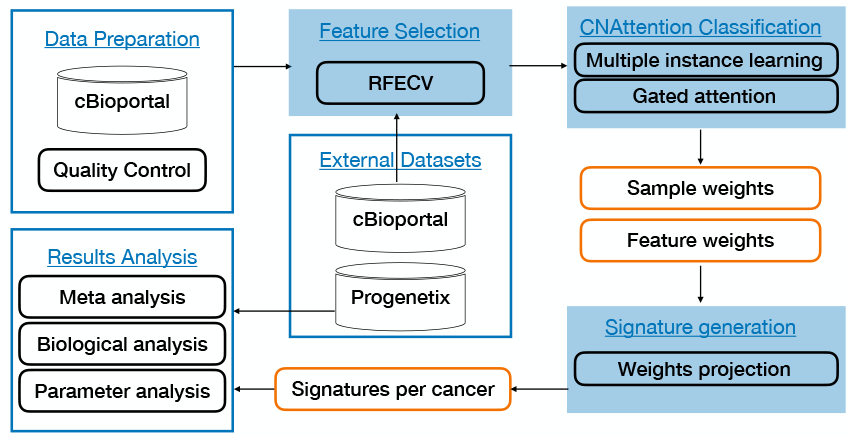
The workflow of the study.

### 2.1 Feature selection by RFECV

RFECV, which stands for Recursive Feature Elimination with Cross-Validation, is a feature selection technique commonly used in machine learning for identifying the most relevant features in a dataset. The RFECV algorithm works by recursively removing features from the dataset and evaluating the performance of the model using cross-validation at each step. It starts with all features and iteratively removes the least important ones based on a specified criterion (e.g., model accuracy or another performance metric).

The RFECV algorithm can be summarized as follows:

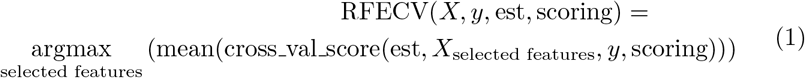

where:

- *X* is the feature matrix,
- *y* is the target vector,
- estimator is the machine learning model used for evaluation,
- scoring is the evaluation metric (e.g., accuracy, precision, recall).

### 2.2 Cancer classification by CNAttention

In typical machine learning problems like image classification, it is assumed that an image clearly represents a category (a class). However, in many real-life applications, multiple instances are observed and only a general statement of the category is given. This scenario is called multiple instance learning (MIL [22, 23]). MIL deals with a bag of instances for which a single class label is assigned. Hence, the main goal of MIL is to learn a model that predicts a bag label, e.g., a medical diagnosis. In our scenario of cancer classification by CNA profiles, since the heterogeneity of cancers, CNA profiles of the same cancer may look different, and we are more focused on the general CNA pattern of cancers. Therefore, we treat each sample as an instance, to ensure that when representative samples of a specific class are in the same bag, this bag is more likely to show a general pattern in this cancer type. In this way, the general pattern of cancers instead of individual CNA profiles are learned. An additional challenge is to discover key instances [24], *i.e.* the instances that trigger the bag label; i.e., the representative CNA patterns of a certain class. By uncovering the key instances, we will be able to reveal the general CNA patterns in the cancer landscape as well as CNA heterogeneity within cancers, which may be related to cancer subtypes. In this paper, we propose a new method CNAttention that extends [25] to multiple classification scenarios of cancer classification based on CNA profiles. We formulate the MIL model using the Bernoulli distribution for the bag label and train it by optimizing the log-likelihood function. We show that the application of the Fundamental Theorem of Symmetric Functions provides a general procedure for modeling the bag label probability (the bag score function) that consists of three steps: (i) a transformation of instances to a low-dimensional embedding, (ii) a permutation-invariant (symmetric) aggregation function, and (iii) a final transformation to the bag probability. We parameterize all transformations using neural networks (i.e., a combination of convolutional and fullyconnected layers), which increases the flexibility of the approach and allows us to train the model in an end to-end manner by optimizing an unconstrained objective function. In addition, we use a trainable weighted average where weights are given by a two-layered neural network. The two-layered neural network corresponds to the attention mechanism [26, 27]. Notably, the attention weights allow us to find key instances.

#### Problem formulation

In the classical (binary) supervised learning problem, the objective is to find a model that predicts a binary value for a target variable, *y* ∈ 0, 1, given an instance **x** *∈* ℝ^*D*^. In the case of the MIL problem, however, instead of a single instance, there is a bag of instances, *X* = **x**_1_, *…*, **x**_*K*_, which exhibit neither dependency nor ordering among each other. It is assumed that *K* could vary for different bags. In our scenario, assume that there are *N* classes (i.e., cancer types), and each instance is assigned a label from 1 to *N*, representing its class or category. The bag label (*Y*) represents the distribution of class labels within the bag. For each class *i*∈ 1, *…, N, Y* [*i*] indicates the proportion of instances in the bag belonging to class *i*. This distribution captures the diversity of class labels within the bag. Furthermore, we assume that individual labels exist for the instances within a bag, i.e., *y*_1_, *…, y_K_*, where *y*_*k*_∈ 1, *…, N* for *k* = 1, *…, K*, however, these labels are not accessible during training and remain unknown. We can express the assumptions of the MIL problem in a concise form using the maximum operator:

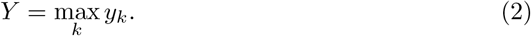

These assumptions indicate that a MIL model must be permutation-invariant. However, optimizing an objective based on the maximum over-instance labels could present challenges, especially in the context of multiple classifications. Gradient-based learning methods may encounter issues such as vanishing gradients, and this formulation is applicable only when employing an instance-level classifier.

To address these challenges inherent in multiple classifications, we train a MIL model by optimizing the log-likelihood function. In this approach, the bag label distribution follows a multinomial distribution, where each class has a probability *θ*_*i*_(*X*) ∈ [0, 1] of being present in the bag of instances *X*. Here, *θ*_*i*_(*X*) represents the probability of class *i* given the bag of instances *X*.

### 2.3 MIL with Neural Networks

In classical MIL problems, it is typically assumed that instances are represented by features that do not require further processing, i.e., *f* serves as the identity function.

However, for certain tasks such as image or text analysis, additional feature extraction steps may be necessary. Moreover, theorems in [28] suggest that for a sufficiently flexible class of functions, any permutation-invariant score function can be modeled. Therefore, we consider a class of transformations parameterized by neural networks *f*_*ψ*_(*·*) with parameters *ψ*, which transform the *k*-th instance into a low-dimensional embedding, *hk* = *fψ*(*x*_*k*_), where *h*_*k*_∈ ℋ and ℋ = [0, 1] for the instance-based approach, and ℋ = ℝ^*M*^for the embedding-based approach.

Ultimately, the parameter *θ*(*X*) is determined by a transformation *g*_*ϕ*_ : ℋ^*K*^→ [0, 1]. In the instance-based approach, the transformation *g*_*ϕ*_ is simply the identity function, while in the embedding-based approach, it could also be parameterized by a neural network with parameters *ϕ*.

The concept of parameterizing all transformations using neural networks is highly attractive, as it allows for arbitrary flexibility in the approach and enables end-to-end training via backpropagation. The only requirement is that the MIL pooling operation must be differentiable.

### 2.4 Attention-based MIL pooling

#### Attention Mechanism

We employ a weighted average of instances (low-dimensional embeddings), where the weights are determined by a neural network. Additionally, these weights must sum to 1 to maintain invariance to the bag size. The weighted average satisfies the conditions of theorem [28], where the weights, along with the embeddings, constitute part of the *f* function. Let *H* = *h*1, *…, hK* denote a bag of *K* embeddings. We apply the following MIL pooling mechanism:

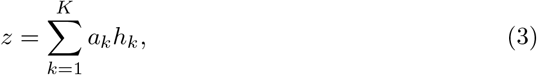

where the attention weights *a*_*k*_ are calculated as follows:

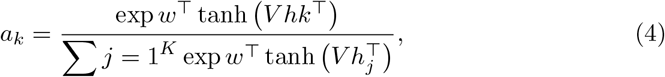

where *w∈* ℝ^*L*×1^and *V ∈* ℝ^*L*×*M*^ are parameters. Additionally, we employ the hyperbolic tangent tanh(*·*) element-wise non-linearity to allow for both negative and positive values, facilitating proper gradient flow. This proposed construction enables the discovery of (dis)similarities among instances.

#### Gated Attention Mechanism

We applied augmenting the tanh(*·*) non-linearity with a gating mechanism, yielding the following formulation:

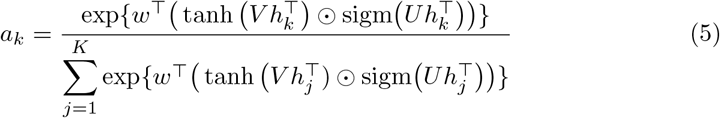

where *w*∈ ℛ^*L*×1^,*V*∈ ℛ^*L*×M^, *U*∈ ℛ ^*L*×M^ are learnable parameters, ⊙ is an element-wise multiplication and *sigm*() is the sigmoid non-linearity. The gating mechanism introduces a learnable non-linearity that potentially removes the troublesome linearity in *tanh*().

The attention-based MIL pooling allows for assigning varying weights to instances within a bag, enabling the bag-level classifier to discern key instances and generate highly informative bag representations. Moreover, coupling the attention-based MIL pooling with transformations *f* and *g* parameterized by neural networks renders the entire model fully differentiable and adaptive. These characteristics render the proposed MIL pooling mechanism highly flexible, capable of modeling an arbitrary permutation-invariant score function.

Ideally, for a bag with the label (*Y* = *i*), high attention weights should be assigned to instances likely to have label *y*_*k*_ = *i* (key instances). Thus, the attention mechanism facilitates easy interpretation of the decision in terms of instance-level labels. While the attention network does not directly provide scores like the instance-based classifier, it can be considered a proxy for it. The attention-based MIL pooling bridges the gap between the instance-level and embedding-level approaches.

### 2.5 Signature generating

After applying MIL, we still need to transfer the bag label to the instance label, therefore transferring the instance weights to feature weights for each unique class. Also, we need the instance label for evaluating our method. To calculate the instance accuracy, we first need to obtain the weighted predictions for each instance. This can be achieved by multiplying the attention weights with the class predictions for each instance within each bag and accumulating these weighted predictions across bags.

Let *w*_*i,j*_ represent the attention weight for instance *i* in bag *j*, and *p*_*i,k*_ denote the class prediction for class *k* for instance *i*. The weighted prediction *wp*_*i,k*_ for class *k* and instance *i* is computed as follows:

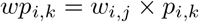

After obtaining the weighted predictions for each instance, we normalize these predictions by dividing them by the sum of attention weights for each instance across all bags. Let *n*_*i*_ represent the normalization factor for instance *i*, which is the sum of attention weights for instance *i* across all bags. The normalized prediction 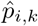 for class *k* and instance *i* is computed as follows:

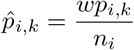

Once we have the normalized predictions for each instance, we can determine the predicted class for each instance by selecting the class with the highest normalized prediction. Let 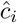 represent the predicted class for instance *i*, which is obtained by:

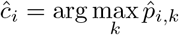

Finally, we compare the predicted classes with the ground truth labels for each instance to compute the instance accuracy, defined as the proportion of instances for which the predicted class matches the ground truth label.

The instance accuracy can be calculated using the following formula:

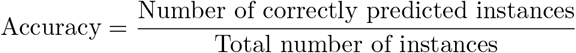

## 3 Results

### 3.1 Datasets

Gene-level CNA profiles across various cancers were obtained from cBioPortal. CNV values were discretized into five categories: “-2” (deep loss, likely homozygous deletion), “-1” (shallow loss, likely heterozygous deletion), “0” (diploid), “1” (low-level gain), and “2” (high-level amplification).

Gene-level CNA profiles of the 30 cancer types (with samples over 50) were provided in the cBioPortal database.The 30 cancer types and the sample numbers are listed in Table 1. After gene feature alignments, there are 10,628 CNA profiles on 24,919 genes. After feature selection by RFECV, the number of gene features dropped to 2917.

**Table 1.**
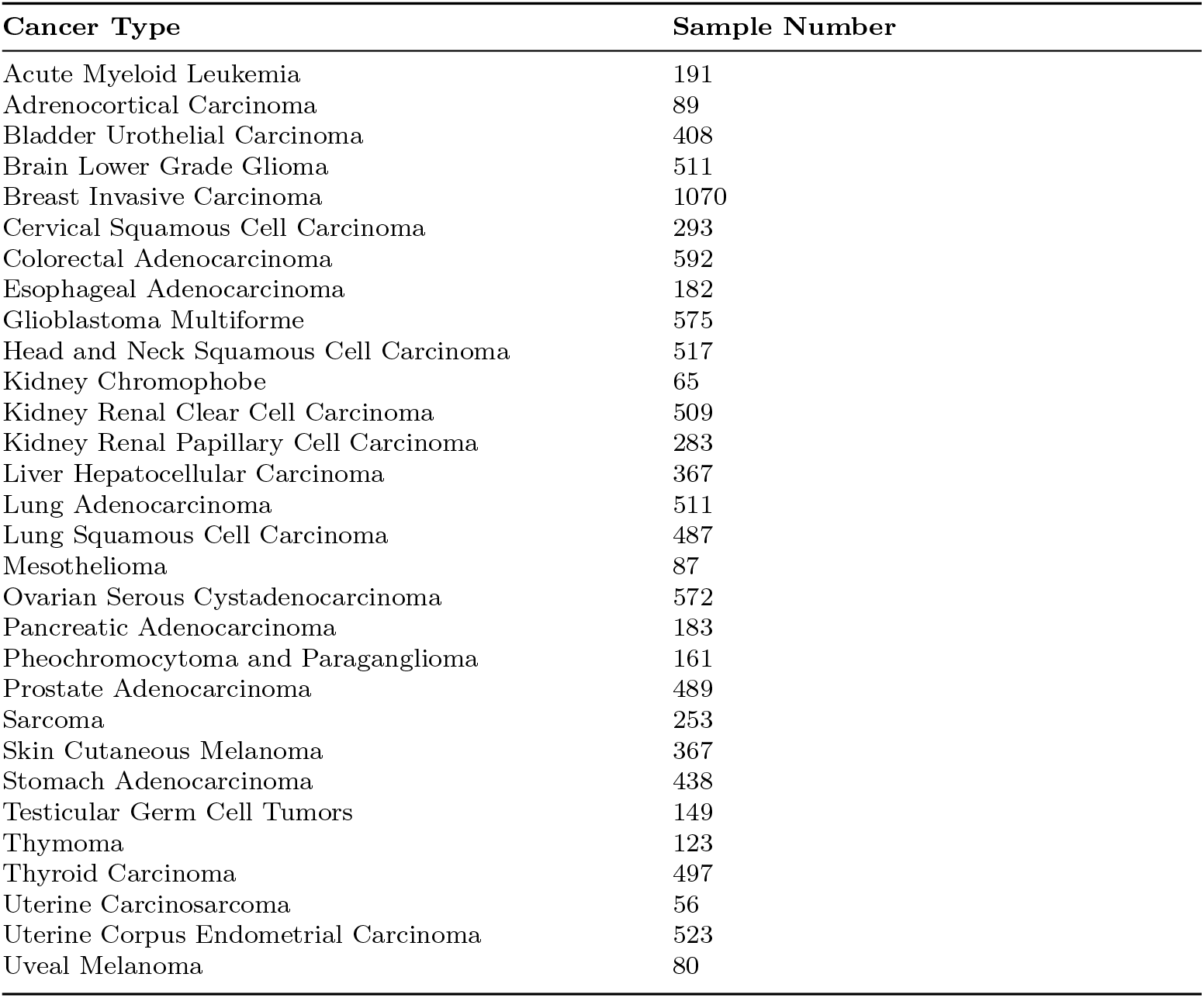
The number of samples and cancer subtypes in the studied cancers.

### 3.2 Classification performance analysis

As shown in Fig. 2, the accuracy for cancer type assignment of samples is high withan average of 0.89. Exceptions are the misclassification of Uterine Carcinosarcoma and Uterine Corpus Endometrial Carcinoma, and Thymoma and Thyroid Carcinoma. A potential reason can be found in Fig. 3 where the entities with lowest scores all fall into area with the lowest sample numbers (below 200). However, for a larger number of entities with less than 200 samples classification accuracy remains sufficient and additional factors such as incorrect diagnostic classification or genomic heterogeneity in those entities might play a role here. For benchmarking of the methodology we compared CNAttention with other methods with and without our feature selection, as shown in Table 2, our method outperforms others. We need to note that comparing to the deep learning method based on keras indicates the effectiveness of the attention mechanism in capturing the CNA patterns of different cancer types. Also, the increase in accuracy of random forest indicates not only the decrease in time costs but also the performance improvement. In addition, we added two external datasets of glioblastoma [29] and colorectal adenocarcinoma [30] for testing, results show that the accuracy remains stable, which proves the robustness of CNAttention.

**Table 2.** Classification comparison with other methods.

| Method | Average accuracy |
| --- | --- |
| **CNAttention** | **0.89** |
| Random Forest | 0.65 |
| Random Forest (without feature selection) | 0.64 |
| [31] | 0.72 |
| [32] | 0.67 |
| Keras [33] | 0.56 |

**Fig. 2.**
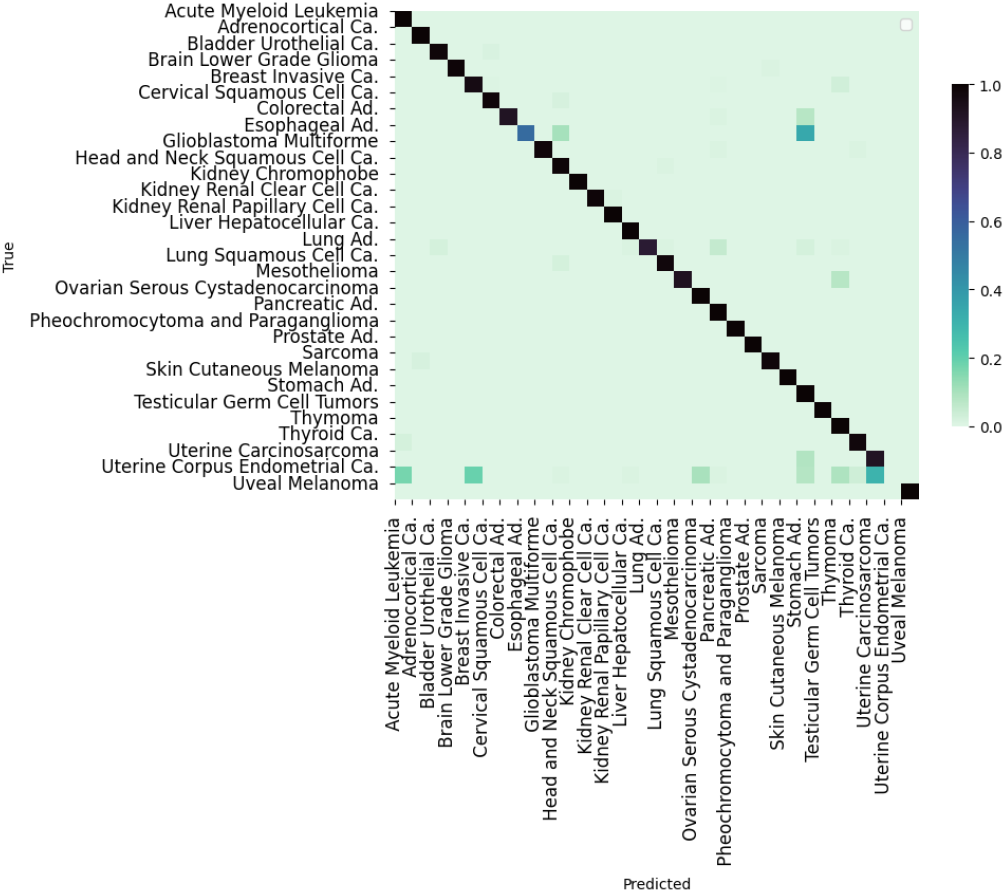
The cancer classification performance of CNAttention. The values in the cells indicate the percentage of biosamples of the corresponding row cancer type classified to the corresponding column cancer type.

**Fig. 3.**
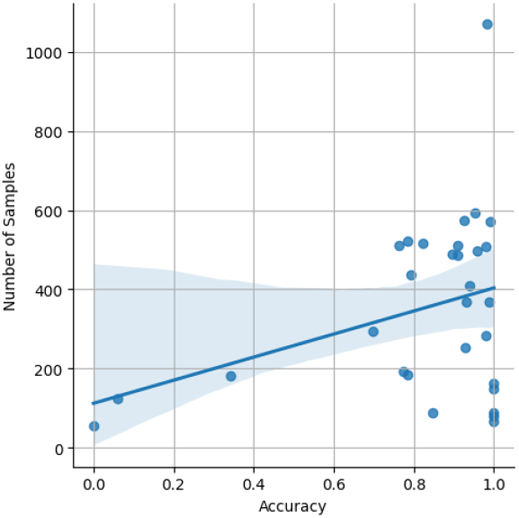
Correlation between classification accuracy and the number of samples

### 3.3 CNA signatures

By using the data processing and modeling procedures described in the Methodology section, we generated a panel of feature genes for each cancer type from the collected CNA samples. Then, with the combination of feature genes of each cancer type, we create an abstract representation for each copy number profile, where only alternations that contributed to the distinctiveness of the sample were preserved. Fig. 4 compares the original CNA patterns with the derived signature features where the frequent and extensive regional alterations in the original data have been replaced by 1008 feature genes, which visibly compare to subsets of characteristic changes in the original CNA data and represent the most discriminative alternations. In addition, we use these features on the random forests model to test whether these genes can represent data without losing key features. The results show that the accuracy remains almost the same, indicating that these features can effectively reduce the dimension of data while maintaining the specific patterns of different cancers.

**Fig. 4.**
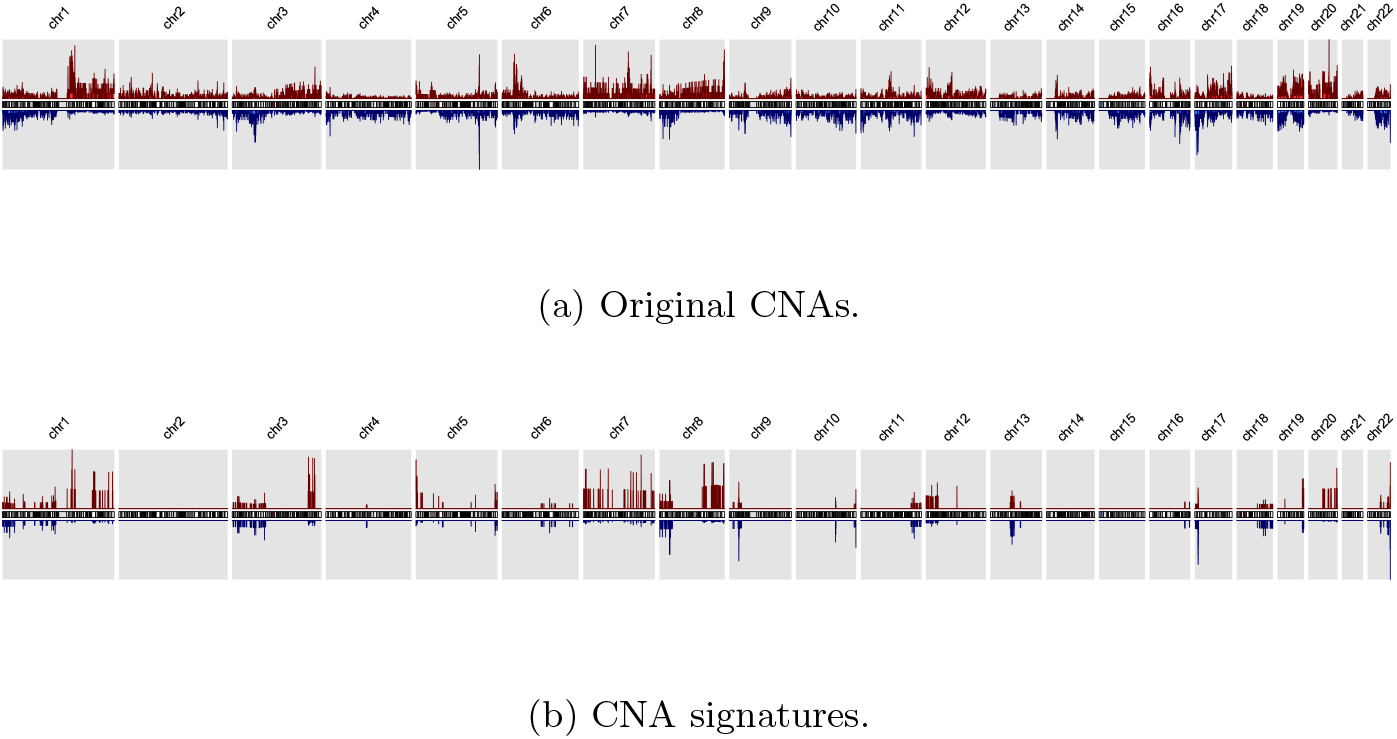
The copy number alternation landscape of all samples using the original CNAs and the feature genes. The feature genes are able to dramatically reduce the complexity of CNA signals while maintaining the mutational characteristics. The red colors above the chromosome axis represent the average amplifications, and the blue colors below the chromosome axis represent the average deletions.

We deduct the set of cancer-related, significantly altered feature genes and could generate the CNA signatures of 30 cancer types, in which these features reflect the sample’s own mutations. Every signature consists of a subset of the general feature genes and their relative signal intensity compared to other signatures. Fig. 5 illustrates a cluster map of the signatures across 30 cancer types. Diseases in the same cluster usually also share close ontologies. Fig. 6 shows the top 30 genes that most frequently occur in signatures, compared to deletions, duplications are more important in cancer development which is consistent that oncogenes are more potent in activation. CDKN2B deletion is the most frequent CNA in all cancer types. We applied them to GO enrichment analysis and results show that they are significant in cancerrelated biological activities including tumor suppressor, human T-cell leukemia virus 1 infection, etc. In the majority of copy number studies, analyses of tumor samples are focused on identifying the driver genes or the focal regions. The genes in the signatures rather reflect the uniqueness of each sample or each cancer type. It is important to note that the feature genes are not intended as the only nor the optimal representation. The fundamental objective of the study is to explore the potential driver mutations that are infrequent but relevant to specific cancer types. Although the signatures do not imply pathogenic causation, we can instead reveal their potential implications and correlations by investigating the signatures and the feature genes. In general, spatial and annotation analysis suggest that some feature genes reflect structural and functional alternations in samples; the signatures of different cancer types show high preference in several genomic regions that suffer frequent CNAs in many cancer types.

**Fig. 5.**
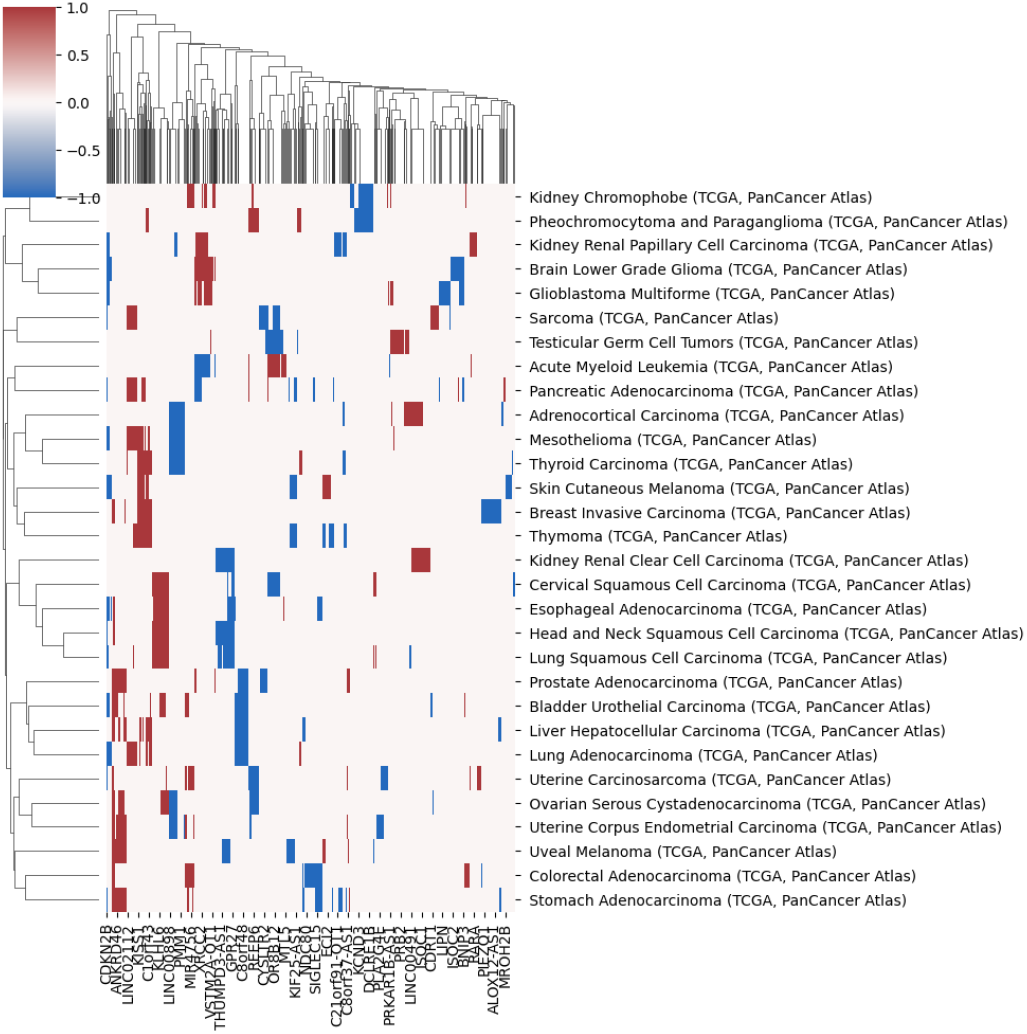
A clustering heatmap of features in 30 signatures. Columns are normalized average CNV intensities of feature genes, where the blue colors are duplication features and red colors are deletion features.

**Fig. 6.**
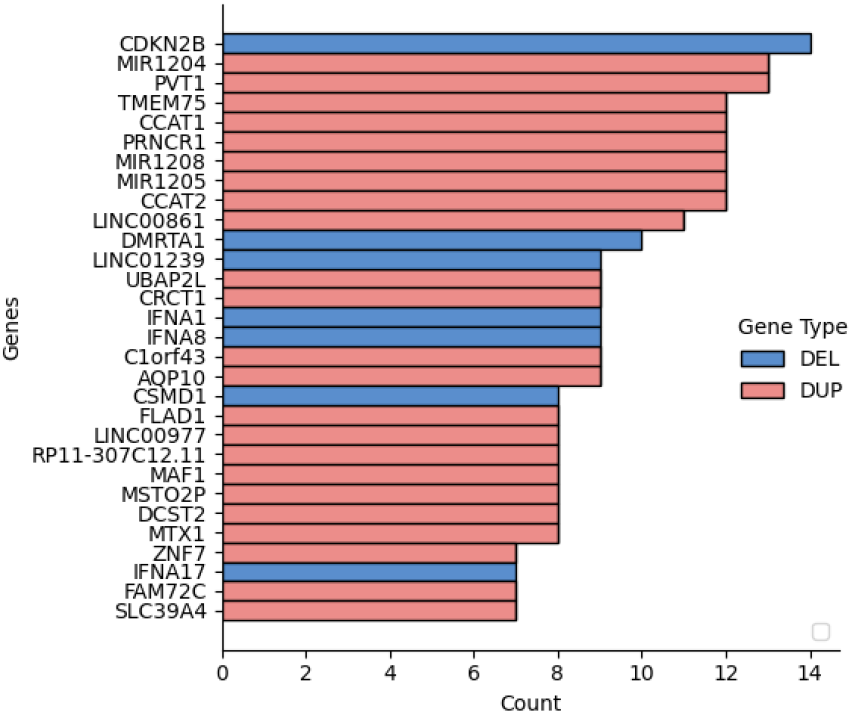
Signature gene count on different cancers.

### 3.4 Validating CNA signatures with a large-scale CNV reference database

We further extend our signatures to external datasets from Progenetix, which includes more heterogeneous CNV profiles from multiple data sources, to evaluate whether our signatures can extract the CNV pattern of certain cancer types. Here are the examples of lung adenocarcinoma and lung squamous cell carcinoma (Fig. 7 and 8).

**Fig. 7.**
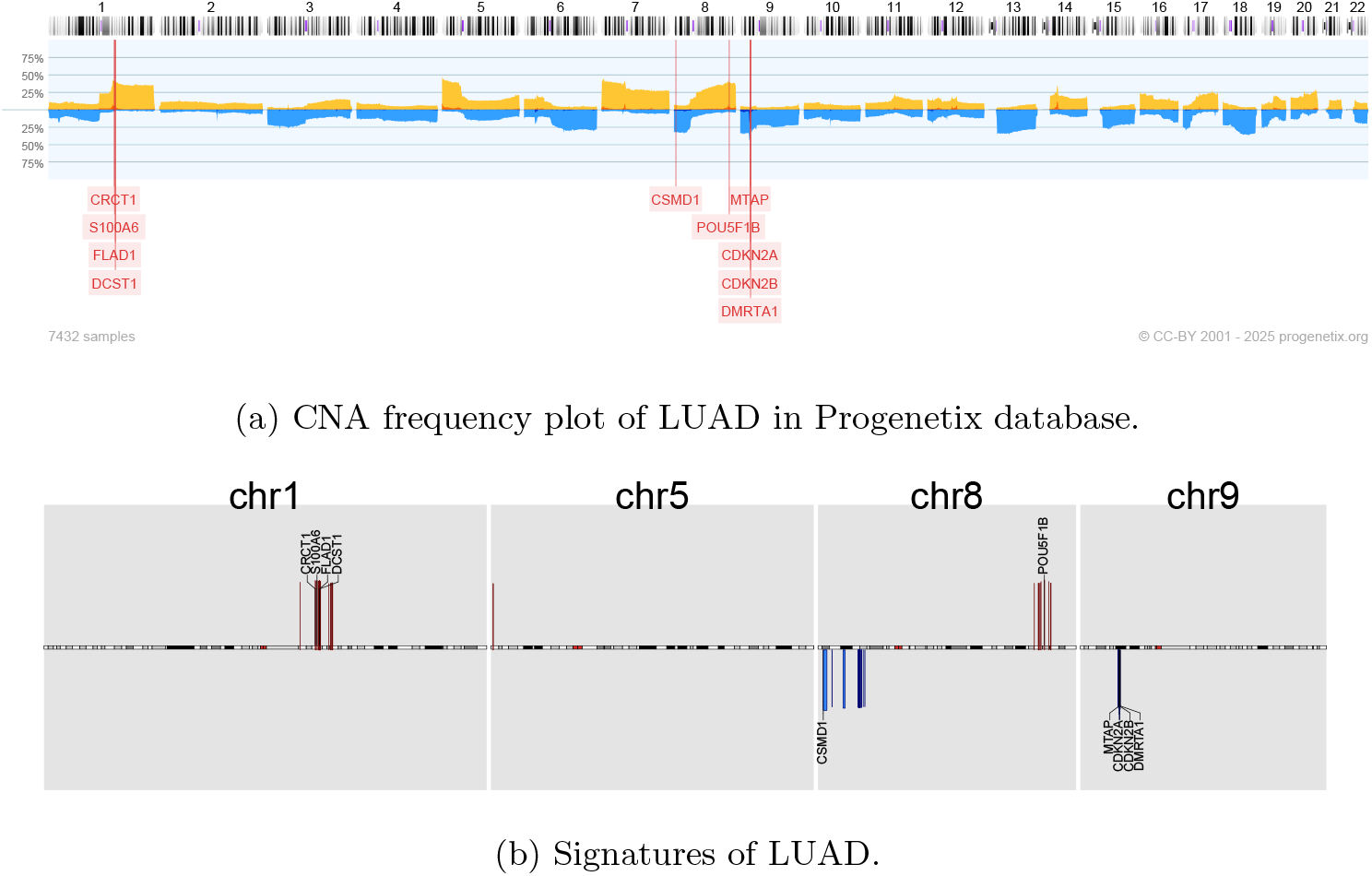
External lung adenocarcinoma CNA frequency plot V.S. gene signatures of lung adenocarcinoma. In the frequency plot, the orange and blue colors indicate duplication and deletion, separately, and the y-axis indicates the frequency. In the signature plot, the red and blue colors indicate duplication and deletion, separately, the y-axis indicates the importance score of the corresponding gene signature.

**Fig. 8.**
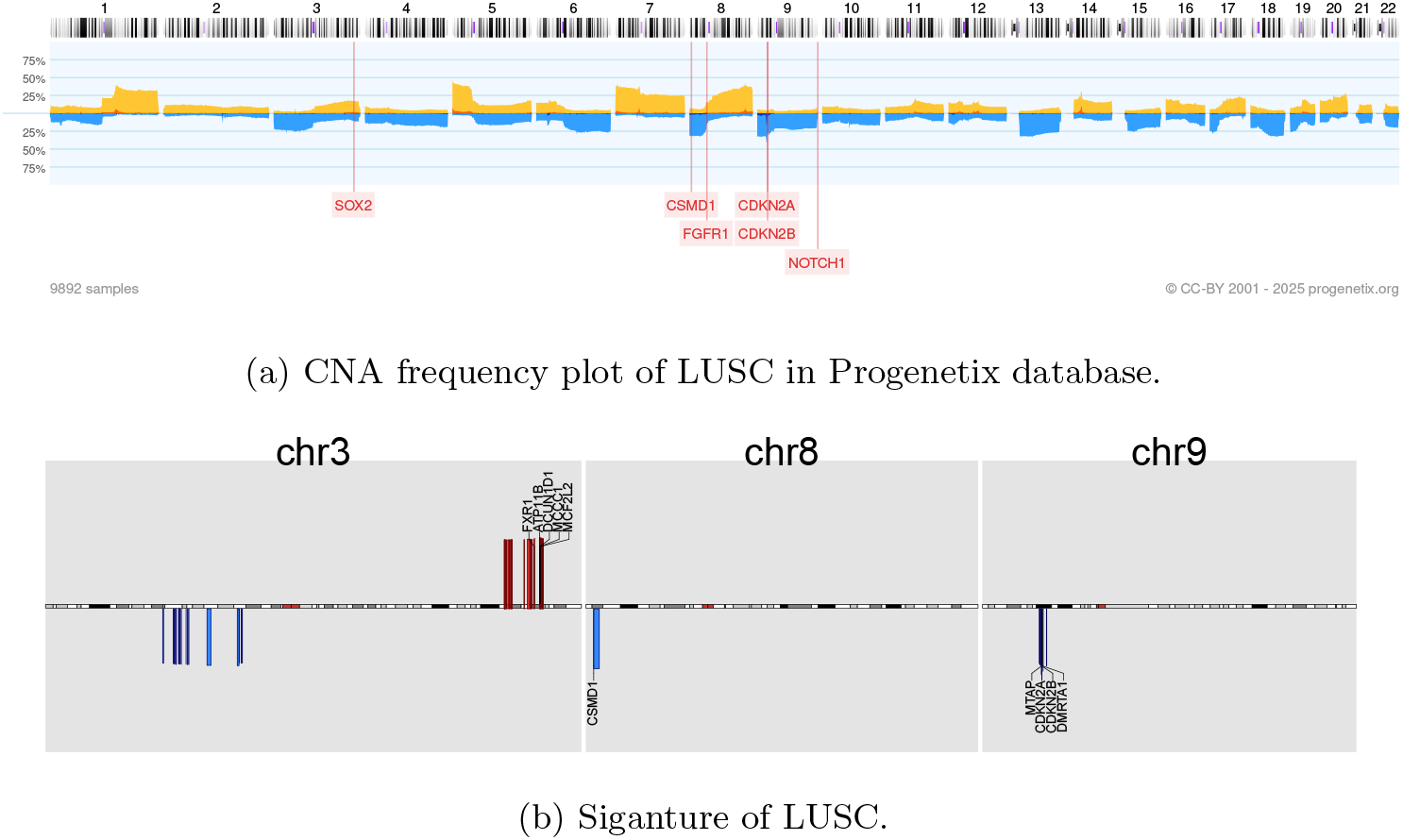
External lung squamous cell carcinoma CNA frequency plot V.S. gene signatures of lung squamous cell carcinoma.

Lung Adenocarcinoma (LUAD) and Lung Squamous Cell Carcinoma (LUSC) represent two major histological subtypes of lung cancer, each characterized by distinct molecular profiles, including unique patterns of copy number aberrations (CNAs). In LUAD, One study has proved that clonal loss of functional TP53 is significantly associated with subclonal gains of MCL-1 (1q21.2) [34]. Distinct CNA patterns characterize LUSC, with frequent amplifications of genes such as SOX2 (SRY-Box Transcription Factor 2) on chromosome 3q26 and FGFR1 (Fibroblast Growth Factor Receptor 1) on chromosome 8p11.23 driving oncogenic signaling pathways essential for cell proliferation and survival. Deletions affecting genes like NOTCH1 (Neurogenic Locus Notch Homolog Protein 1) on chromosome 9q34.3 are commonly observed, leading to dysregulated signaling cascades and accelerated tumor progression. In addition, loss of tumor suppressor genes CDKN2A/2B and CSMD1 are shared between LUAD and LUSC signatures. We can conclude that signatures can extract the features with relevance to the specific cancer, and also keep the common CNA pattern between LUAD and LUSC, help uncovering the relationships between different cancers.

### 3.5 Similarities of Neural Crest Originated Subtypes

We further extend the signatures to more external datasets, by importing the cancer classification tree in Progenetix, and we find the similarities of neural crest-originated subtypes, including glioblastoma, glioma, medulloblastoma, and melanoma. The four distant cancer types exhibit highly similar signatures in both feature selection and alteration frequencies. Figure 9 illustrates the comparison of original CNA data, features, and known drivers of these cancers on chromosomes with similar signatures. Notably, their signatures show high similarities in the duplication of chromosome 7 and the deletion of chromosomes 9 and 10. Additionally, they share pairwise similarities in the duplication of chromosome 1 and 20, as well as the deletion of chromosome 14.

**Fig. 9.**
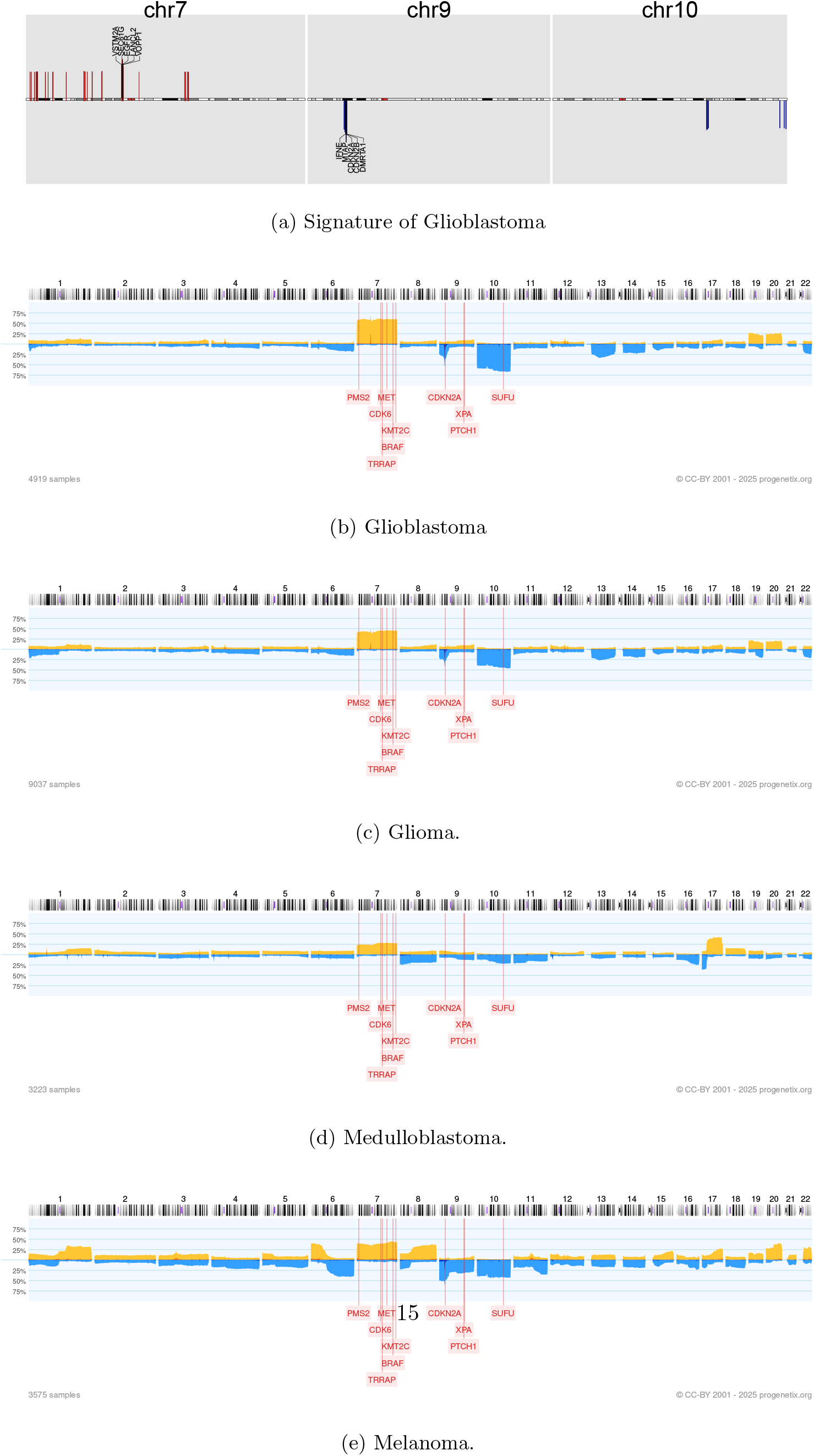
CNV signature of Glioblastoma V.S. CNV frequency of Glioblastoma, Glioma, Medulloblastoma and Melanoma in Progenetix database. These neural crest originated subtypes show similar CNV patterns

Chromosome 7, with frequent copy number gains in all four cancers, harbors several key oncogenes such as EGFR, CDK6, and MET in glioma; KMT2C and PMS2 in medulloblastoma; BRAF, RAC1, and TRRAP in melanoma. Similarly, chromosome 9 and 10, commonly deleted in these cancers, contain several important suppressor genes such as CDKN2A and PTEN in glioma; XPA, PPP6C, and CDKNA in melanoma; PTCH1 and SUFU in medulloblastoma. Notably, the CDKN2A/B deletion is the most frequent copy number aberration across all cancer types.

In the 1990s, epidemiological studies [35] initially uncovered a link between melanoma and nervous system tumors. This association was not only observed in familial cases, confirmed by germline mutations, but also indicated a significantly elevated risk of one disease in individuals with a history of the other. Despite evidence suggesting shared pathophysiological pathways and responsiveness to similar drugs, the genetic connection between these two disease groups remained largely elusive [36]. Despite their clinical and histological disparities, medulloblastomas, melanomas, and gliomas all originate from neural crest cell lineages. Recent research on neural crest cells and cancers derived from these lineages suggests that malignant cells mimic various aspects of neural crest development at a behavioral, molecular, and morphological level [37]. Aberrations in tumor cells may reactivate embryonic developmental programs, thereby promoting tumorigenesis and metastasis. In melanoma, WNT family members, crucial during the epithelial-to-mesenchymal transition of neural crest cells, are reactivated during invasive transformation [38]. In glioblastoma, experimental data suggests dysregulated WNT signaling supports the onset of cancer stem cells, facilitating tumor enlargement and metastasis [38]. In medulloblastoma, the WNT subgroup represents one of the four molecular subtypes of the disease [39]. Regions with similar signatures show abnormal amplification frequencies in several WNT genes, potentially reflecting overexpression of WNT signaling. Specifically, WNT2B and WNT4 exhibit moderate amplification frequencies, while WNT2, WNT3A, WNT9A, and WNT16 demonstrate high amplification frequencies. WNT2 and WNT16 are signature genes in all three subtypes, indicating their prevalence among individual samples.

### 3.6 CNV heterogeneity reflects cancer subtypes

To find out whether these signatures reflect the heterogeneity within cancers, we collected all available clinical information and adopted a random forest to see whether the signatures could help classify the subtypes. The results show that for all cancers, compared to using all CNV profiles, using signatures can help increase the accuracy, indicating that signatures can reflect the heterogeneity of subtypes within cancer. Here we showcase how signature reflects subtypes of Brain Lower Grade Glioma in Fig. 10 by the 1p/19q co-deletion.

**Fig. 10.**
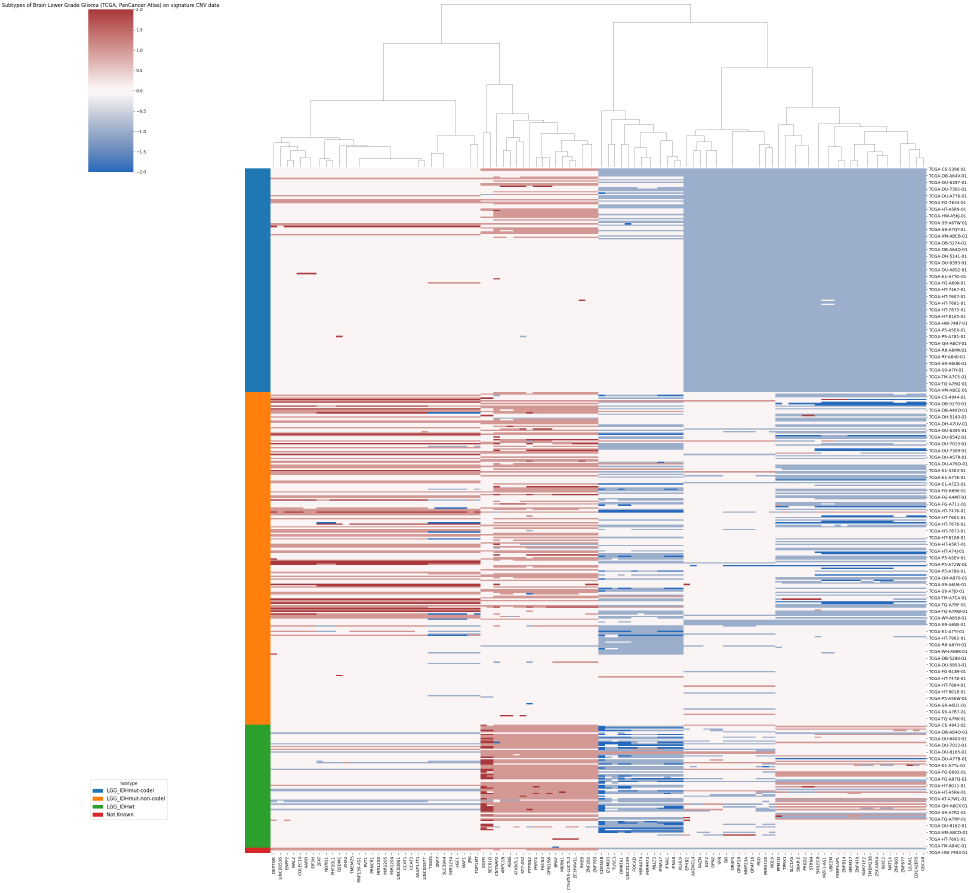
Subtypes of Brain Lower Grade Glioma on signature CNV data. The different subtypes show three different CNA patterns.

## 4 Discussion

In this study, we compiled an extensive collection of cancer Copy Number Alteration (CNA) profiles to pinpoint genomic aberration signatures specific to each cancer type. Our novel attention-based MIL method, CNAttention, showcased the potential of CNA patterns in tumor identification. Focusing on isolating unique components in diagnostic CNA profiles, we extracted signatures for 30 cancer types, each characterized by a minimal gene representation with high discrimination capacity.

Comparative analyses of signature genes and their respective regions revealed frequent duplications on chromosomes 7 and 8, and deletions on chromosome 22 across various cancer types. Despite their prevalence, these regions also exhibited features with high differentiating power, possibly indicating the functional significance of cancer-related genes specific to pathway involvement.

Our analyses unveiled shared CNA signatures among four clinically and pathologically distinct cancer types—glioblastoma, medulloblastoma, melanoma, and glioma. These tumor types can be traced back developmentally to common lineages of neural crest cells. Although research has sporadically linked neural crest cells with glioma and melanoma development, the genetic underpinnings of their association in oncogenetic processes remain elusive. Through comparative analysis of shared mutations and improvements in developmental processes in normal tissues, insights into shared pathologies and potential therapeutic targets may emerge. In addition, the signatures reveal the heterogeneity of cancer types, and shed light on uncovering more potential cancer subtypes.

In summary, this study presents a systematic pipeline for integrative and comparative analyses of a large amount of copy number data. The resulting CNA signatures offer new perspectives on the understanding of common foundations in cancers and show promising potential in applications of tumor classification.

